# Selection for longevity is stronger in larger populations but not in more stable populations

**DOI:** 10.64898/2026.11.21.765981

**Authors:** Caroline E Freeland, Kailey R Muncey, Lev Y Yampolsky

## Abstract

Many features of biology, including life-history, are explained by either neutralist or adaptationist arguments. Concordantly, two explanations have been proposed for inter- and intraspecific variation in lifespan in *Daphnia*. Fitness and longevity correlate with local population size, consistent with accumulation of deleterious mutations in small populations (Lohr & Haag 2015). On the other hand, fitness and longevity are lower in species and genotypes evolving in intermittent habitats, implying selection favoring reproduction at the expense of longevity (Dudycha & Tessier 1999). Here differentiate between the two explanations by measuring lifespan and reproduction investment in 33 clones of *D. magna* originating from a habitats of different size and. We found a correlation between median lifespan and size of the habitat of origin, implying the effect of drift in small populations, but no differences among habitat types (permanent vs. large intermittent), i.e., no indication of stronger selection for longevity in permanent habitats. We found positive correlations between longevity and offspring quality at both *ad libitum* and restricted food levels, and between longevity and offspring number at *ad libitum* food level, consistent with the genetic load explanation. However, a negative correlation between longevity and fecundity was observed when food was limited, indicating longevity-fecundity trade-off.

## Introduction

Life-history traits are shaped by multiple evolutionary forces, and their interpretation often reflects a long-standing debate between adaptationist and neutralist perspectives in evolutionary biology. Lifespan and senescence may evolve through adaptive optimization under local ecological conditions but can also emerge as a consequence of mutation accumulation and imperfect purifying selection, representing a classic selectionist/neutralist argument (Medawar 1952; Williams 1957; Hamilton 1966; Charlesworth 2000; Li et al. 2023). The relative contribution of the two processes – deleterious mutations accumulation due to either small population size or late-age effects and fixation of beneficial alleles through the action of trade-offs between longevity and other life history parameters – is still a matter of debate.

Furthermore, trade-offs between longevity and early reproduction can shape selection differently depending on stability of the environment, with selection favoring early reproduction (or univoltine life cycle, or semelparity) over longevity in short-lived, intermittent, or risky habitats and longevity (perennial life cycle, iteroparity) over early reproduction in more stable habitats (Charlesworth 1994; Dudycha & Tessier 1999; Terzibasi Tozzini & Cellerino 2020; Fisher & Salguero-Gómez 2026)

Perhaps paradoxically, both Medawar’s mutation accumulation theory and Williams’ negative pleiotropy theory predict shorter lifespan in short-lived, risky environment, albeit for different reasons. The essentially neutralist Mutation accumulation theory (Medawar 1952) predicts shorter lifespan in short-lived habitats because the old age “selection shadow” is more radical in the environments where no individuals can possibly live longer than the lifespan of an intermittent, seasonal habitat. In adaptationist models, lifespan evolves as part of a life-history strategy balancing investment into somatic maintenance and reproduction. Classical evolutionary theories of aging predict genetic (Negative Pleiotropy) or resource-focused (Disposable Soma) trade-offs between early-life reproductive output and late-life survival, such that increased allocation to reproduction may come at the expense of longevity (Williams 1957; Kirkwood & Rose 1991; Kowald & Kirkwood 2015). Because environmental predictability and mortality regime are expected to shape this balance, it is reasonable to assume that selection favoring early reproduction at the expense of longevity and somatic maintenance is stronger in temporary or intermittent habitats, where long-term survival and, thus, future reproductive opportunities are uncertain or unlikely (Charlesworth 1994; Roff 2002; Lind et al. 2026). Not surprisingly, examples of short lifespan in intermittent, seasonal, or otherwise predictably risky environments are abundant. The very existence of annual plants and univoltine insects in highly seasonal temperate, polar and, in particular, arid environments are prime examples (Friedman 2020; Numata & Shintani 2023; Poppenwimer et al. 2023), but there are many others across a variety of taxonomic groups (Varpe 2017; Zák, et al. 2021; Lind et al. 2026).

On the other hand, shorter lifespan (and, conceivably, lower reproduction output) may be caused by deleterious mutations escaping stabilizing selection. There are many scenarios in which this is possible, but in the context of life history evolution two are most commonly invoked: accumulation of deleterious mutations with late-age effects, in populations of any size (Medawar 1952), and accumulation of deleterious mutations with age-independent effects in small populations (Kimura 1962). Both scenarios render stabilizing selection inefficient. A trade-off with early fitness and/or investment into reproduction is not required, in fact one might expect a positive correlation between reproduction and longevity, if deleterious mutations affect both. Accumulation of harmful alleles affecting physiological performance and survival in small populations is well documented in various organisms (Cui et al. 2019; Willemsen et al. 2020; Aubier & Galipaud 2023).

The plankton crustacean *Daphnia* is particularly suitable model to differentiate between the two lines of argument, because it inhabits a broad range of habitats from rock pulls less than 1 m^2^ in area and less than 1 m^3^ in volume that contain liquid water only weeks each year to large permanent ponds and lakes. Consequently, both lines of argument have gained support from *Daphnia* dat. Consistent with the view that short lived habitats should promote reproduction over longevity, genotypes and species of *Daphnia* originating from intermittent and ephemeral habitats have been reported to exhibit shorter lifespans and altered life-history allocation compared with those from more stable environments (Dudycha 2003; 2004; Dudycha & Tessier 1999; Dudycha & Hassel 2013), in a series of comparisons involving sister species *D. pulex* and *D. pulicaria*. In contrast, in an interspecific, across population comparisons, longevity and several fitness-related traits have been shown to correlate with local population size, and populations’ mean heterozygosity, supporting the possibility that mutational load contributes substantially to lifespan evolution (Lohr et al. 2014; Lohr & Haag 2015). Thus, habitat size rather than habitat stability may shape interspecific variation in lifespan and other fitness-related traits through different levels of drift and inbreeding in populations of different size.

Distinguishing between these alternatives is difficult because habitat permanence and population size are often confounded in natural populations. Small rockpools and ponds are frequently both intermittent and demographically small, whereas permanent lakes are typically both stable and large. Consequently, ecological selection for shortened lifespan in unstable environments and mutation accumulation in small populations may produce superficially similar predictions. However, some *Daphnia* populations inhabit large but intermittent water bodies, such as summer-dry ponds in Mediterranean climates (Zacharias et al. 2007), small but permanent ponds in temperate climates, or rockpools that are both small and short-lived due to drying in the summer and freezing to the bottom in winter (Jocque et al. 2010). This provides an opportunity to at least partially separate the effects of habitat stability from those of population size. Here, we test competing explanations for variation in lifespan by measuring longevity in 33 clones of *Daphnia magna* originating from habitats differing in size and permanence across the western Eurasian clade’s range. Specifically, we ask whether lifespan is better predicted by habitat stability, consistent with life-history trade-offs and adaptive optimization, or by habitat size, consistent with mutation accumulation in populations experiencing stronger genetic drift. We then ask whether a reproduction vs. longevity trade-off exist and if yes, how is it modulated across habitats of different size and stability.

In summary, the predictions of the two models are as follows. The adaptation to intermittent habitats model predicts shorter lifespan in intermittent habitats, with a trade-off (negative correlation) between lifespan and early reproduction investments. The neutralist model predicts shorter lifespan in populations originating from smaller habitats, regardless of their stability, with non-negative correlations between lifespan and reproduction. We present here an experimental test of these predictions using 33 clones of *Daphnia magna* Straus selected to represent both a range of genetically determined lifespan and stability and size of their habitat of origin, with the size of the habitat, ranging from meters to kilometers, serving as a proxy for population sizes.

## Methods

### Clones’ provenance and maintenance

All clones of *D. magna* used in this study were obtained from *Daphnia* stock collection at Basel University (Switzerland) where they had been maintained for several years after sampling from nature. Clones were selected to represent the entire range of the W.Eurasian clade of *D.magna* (Supplementary Fig. S1) and a broad range of habitat types. Supplementary Table S1 presents data on clones’ origin, including size and stability (in terms of suitability for *Daphnia* existance in the plankton phase) of the pond or lake from which they were originally sampled. Clones were maintained in the lab through parthenogenetic reproduction under 12:12 photoperiod in ADaM reconstructed pond water medium (Klüttgen et al. 1994), fed with *Scenedesmus* green alga culture.

### Lifespan experiment, fecundity, offspring size and lipid content

Prior to the experiment a single founder female was randomly selected from the stock of each clone and propagated asexually. Offspring of these females were used as mothers of experimental animals. They were maintained in groups of 5 in 100 mL jars in ADaM medium, fed by adding *Scenedesmus* culture daily to the concentration of 2E5 cells/mL. Their offspring from 2^nd^ to 4^th^ clutch were collected within 24 hours after birth and transferred initially into the same kind of jars and then, at the age of 6-7 days into lifespan experiment trays.

The lifespan experiment was set up using plastic inserts as described in Beam et al. (2024). Briefly, plastic inserts made out of 50 mL conical centrifuge vials with cut-off bottoms replaced by 1 mm nylon mesh, each containing 5 adult *Daphnia* from the same clone, were placed into 4x8=32 slot racks, which were, in turn, placed into 3 L polycarbonate plastic containers filled with ADaM medium matching the cohort size to maintain the density of 1 adult *Daphnia* per 20 mL. Food was added daily to the concentration of either 1E5 (low food) or 4E5 cells/mL (high food) with 4 replicated containers in each food level. The inserts were lifted from the containers twice daily to remove neonates and retain adults. The census was conducted every other day and the medium was replaced every 4 days, with medium volume adjusted to account for cohorts’ attrition. At each census not coinciding with medium replacement, medium from all containers with the same food concentration was mixed together and returned to the containers. Thus, *Daphnia* from all clones shared the environment constantly with each replicate container and were exposed to the same medium periodically across the food level, ensuring uniform medium composition and microbiome.

At the cohort ages of 21.8 ±3.0 days, 91.5 ±3.1 days and 134.0 ±2.7 days (hereafter “young”, “old” and “very old” samples) a subset of females was sampled from the cohorts, with each females placed into a separate vial containing 20 mL of ADaM medium with food level matching that in the cohorts. Offspring of these female born within a 72 hours were counted and 1-3 neonate per clutch stained with Nile Red (1 mg/mL) for 1 h. Females were then returned to their cohort containers. After staining the neonates were photographed individually under a 10× objective lens with 0.25 aperture on a Nikon Labophot-2 microscope equipped with EFD-2 epifluorescence block and a Motic M5 color camera, with a 510-560 nm excitation and DM 575 nm, BA 590 nm emission filter. Images were analyzed using ImageJ (Rasband 2018) with the portion of pixels with intensity over a threshold recorded as the measure of lipid content (Supplementary Fig. S2A). The threshold was selected separately for each series of images (maternal age), as different series differed slightly in intensity histogram shape. Because of that fluorescence above threshold values were d to each series maximum and no comparisons of neonate lipid content between mothers of different ages were made; a common garden experiment in which offspring of mothers of different ages were measured together will be published elsewhere. The same individuals were also photographed in brightfield to measure body length (from the top of the head through the dorsal edge of the eye to the most distant point on the posterior edge of the carapace, Supplementary Fig. 2B). One clone did not survive to be included into the “old” samples, and two additional clones did not survive to be included into the high food “old” samples. Four clones did not survive to be included into any of the “very old” samples, nine additional clones did not survive to be included into the low food “very old” samples and four additional clones did not survive to be included into the high food “very old” samples.

### Statistical analysis

Using a single clone from a pond of a given size maximized the number of pond sizes we could include, but because each clone was associated with a single pond size, clone identity and pond size were completely confounded. Consequently, any effect attributed to pond size represents a between-clone association and cannot be cleanly separated from other clone-specific differences. Therefore, we conducted three lines of survival analysis. First, we analyzed the effects of food concentration and clones, both as fixed effects, on baseline survival and aging rate using Parametric survival model with Weibull distribution. In this and all further analyses Weibull distribution was found to be the best fit for the data, followed by log-logistic distribution, by Akaike Information Criterion (Supplementary Table S2) and both parameters of Weibull distribution were fitted to the model’s effects. We then implemented the same model with food, habitat type (permanent vs. intermittent vs. intermittent rock pool; a categorical effect), and pool size (log-transformed; a continuous effect), but without clones in the model. These two analyses were carried out using Survival platform of JMP Ver. 19 (JMP 2025). Finally, we then included clones as a random nested frailty effect using Royston–Parmar flexible parametric survival model implemented in the R package rstpm2 (Liu et al. 2018). We also analyzed the data using a Proportional Hazard approach both with and without clones as a random factor, but, because Parametric survival models found a significant change of hazard with age, the proportional hazard models were deemed insufficient for the data and these results, superficially similar, are not reported.

Clone- and food-specific parameters of Weibull distribution, namely “location” parameter interpretable as baseline survival (or baseline longevity) and “delta” interpretable as inverse of age-specific mortality (or of actuarial aging rate) were then correlated with habitat parameters and other clone- and food-specific measurements of life-history traits. These two parameters will be referred to as longevity and aging parameters, respectively. Because the parameters are inherently inversely related due to the properties of fitted parametric model, the residuals from the linear regression of the aging parameter on the longevity parameter were used in downstream analysis instead of raw values, to analyze the correlations of the aging parameters independently. Because two of the clones had unusual short lifespans, Cauchy robust regression (JMP 2025) was used in addition to regular linear regression to capture relationships not caused exclusively by these two outliers.

## Results

### Correlations between parameters of the habitat of origin and longevity

Survival curved of the 33 clones at two food levels (individually and combined by habitat type) are presented on Fig. 1. Food level had a significant effect on both longevity and aging parameters of Parametric Survival Weibull distribution (Table 1), with restricted food resulting in lower baseline survival with longevity parameter of -0.0349 (95% CI -0.0580..-0.0119) but also in lower aging rate (aging parameter 0.0794 (95% CI 0.0092..0.097). This result is corroborated by the Royston–Parmar analysis detecting a highly significant Food-by-Age interaction (P = 2.4e-11; Supplementary Table S3). Likewise, both Clones and Clone-by-Food interactions were significant effects for both parameters. On the other hand, only the effect of Food level on the aging parameter, but not on baseline longevity remained significant in the model in which clones were pooled and habitat stability and size used as fixed effects (Table 2, Fig. 2). This analysis also revealed that pool size was the strongest predictor of longevity, along with pool size-by-habitat type interaction. In contrast, pool size had no effect on aging rate parameter, but habitat type did, with *Daphnia* large intermittent ponds showing slower aging than those from large permanent ponds (contrary to the prediction based on adaptation to habitat stability), while *Daphnia* from intermittent rock pools showing faster aging than those from permanent ponds (Fig. 2B). As expected, inclusion of clones as a random frailty effect in the Parametric Survival model yields a highly significant clone effect rendering nearly all other effects non-significant (Supplementary Table S3), with the exception of the rockpool vs. permanent pond contrast (P<0.02). Thus, overall effects of habitat properties on longevity parameters are hard to untangle from the variance among clones and therefore downstream analysis will focus on clone-specific parameters rather than on effects of clones pooled by habitat type or size.

**Figure 1.**
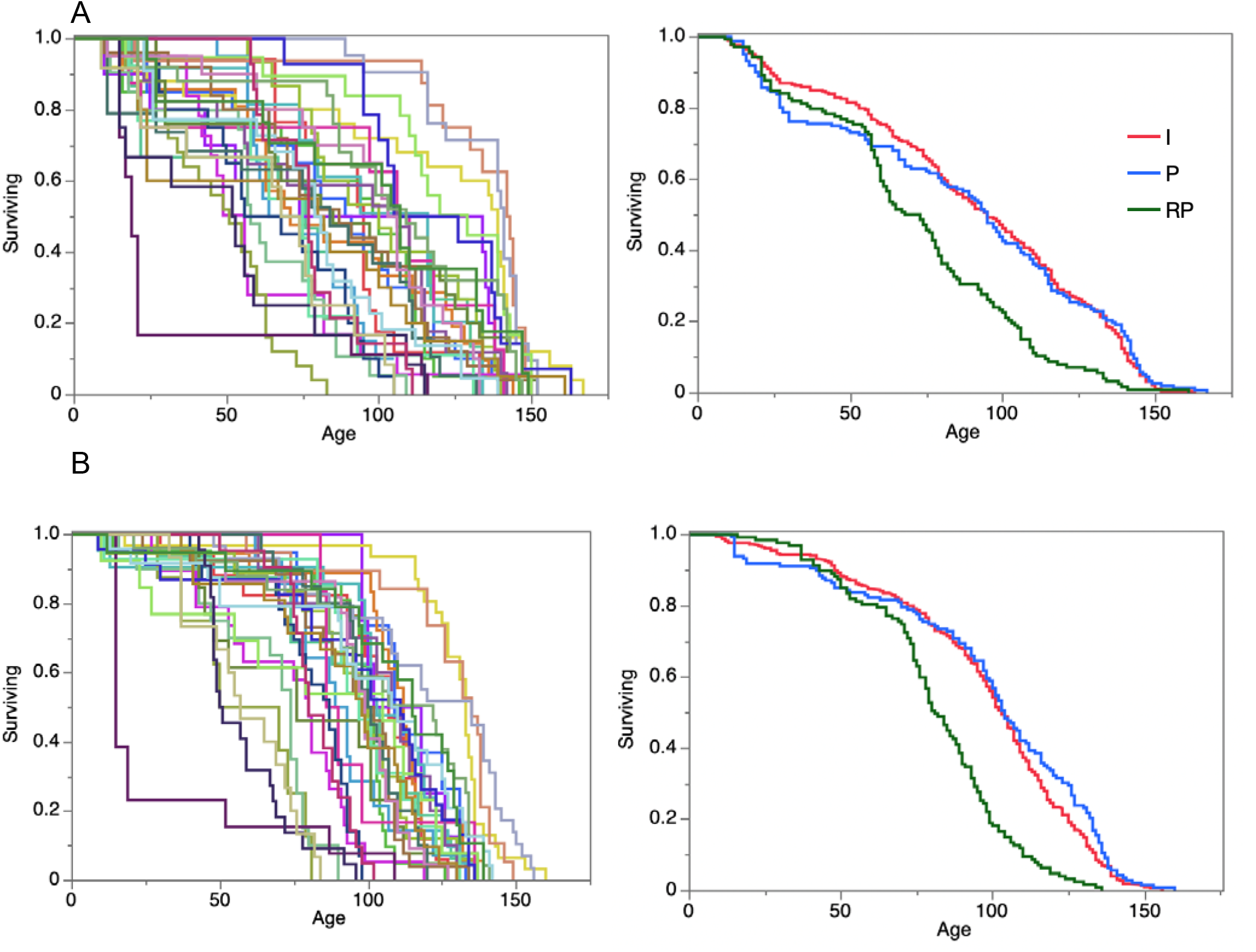
Survival curves of 33 clones (left) and clones combined by habitat type (I=Intermittent, P=Permanent, RP=Rock pools). A: Food level 1e5 cells/mL/day, B: Food: 4e5 cells/mL/day.

**Fig. 2.**
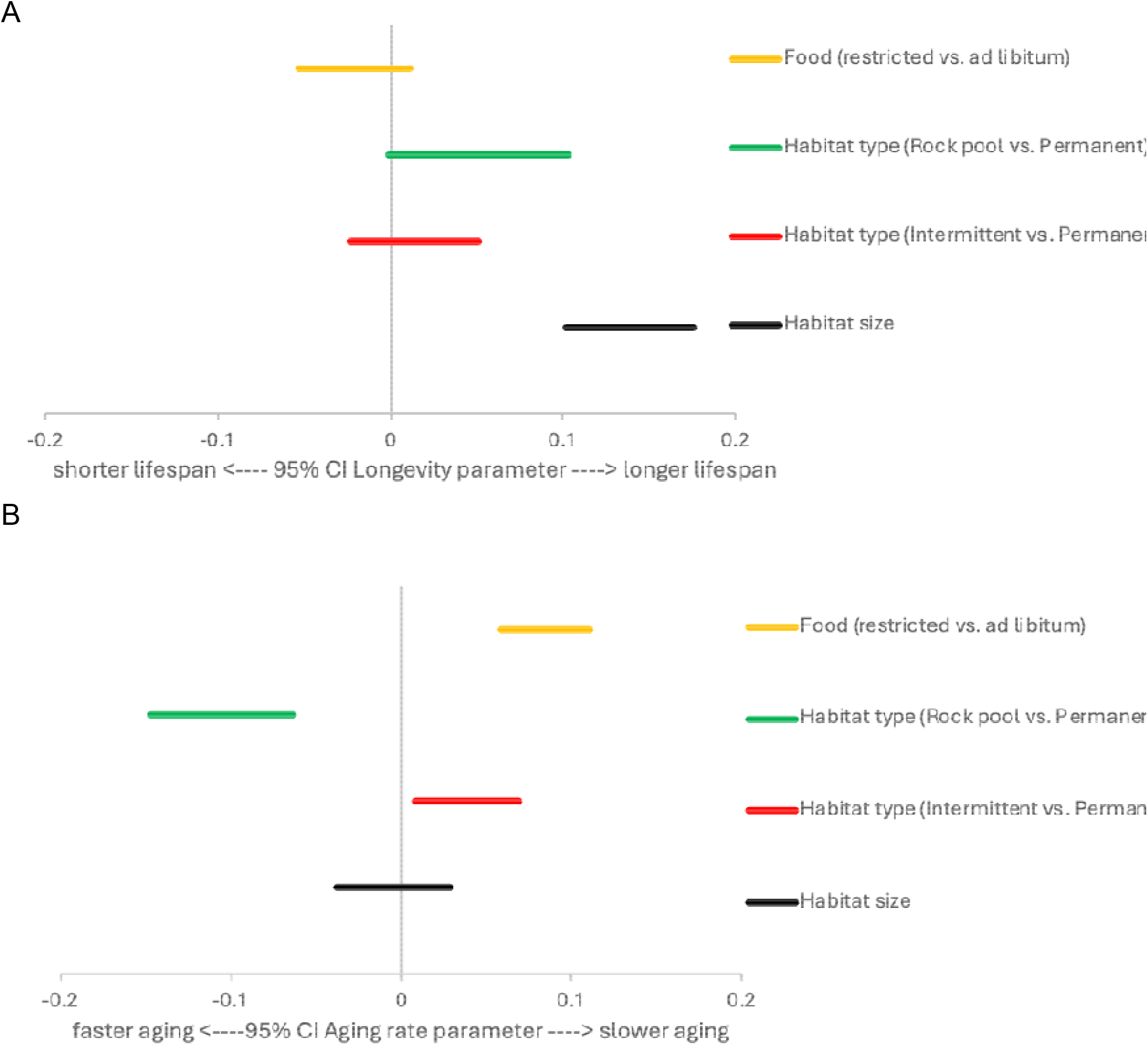
Forest plots of 95% CI’s of the effects of food level and habitat of origin type and size on longevity (A) and aging parameters in Parametric Survival analysis.

**Table 1.** Results of Parametric Survival analysis of longevity of 33 *D. magna* clones under 2 food regimens. See Supplementary Table S1_ModelsComprison for comparisons of different assumed distributions.

| Source | DF | L-RChiSquare | Prob>ChiSq |
|---|---|---|---|
| **Longevity parameter** |  |  |  |
| Food | 1 | 8.745 | 0.0031 |
| Clone | 32 | 314.449 | <0.0001 |
| Food*Clone | 32 | 47.211 | 0.041 |
| **Aging rate parameter** |  |  |  |
| Food | 1 | 44.350 | <0.0001 |
| Clone | 32 | 196.900 | <0.0001 |
| Food*Clone | 32 | 124.958 | <0.0001 |

**Table 2.** Results of Parametric Survival analysis of longevity of *D. magna* from originating from habitats of different type (permanent vs. intermittent vs. intermittent rock pools) and size under 2 food regimens.

| Source | DF | L-RChiSquare | Prob>ChiSq |
|---|---|---|---|
| **Longevity parameter** |  |  |  |
| Food | 1 | 1.632 | 0.20 |
| Habitat(P/I/rp) | 2 | 7.777 | 0.021 |
| Food*Habitat(P/I/rp) | 2 | 4.327 | 0.11 |
| Log<sub>10</sub>(PoolLength) | 1 | 49.913 | **<0.0001** |
| Food*Log<sub>10</sub>(PoolLength) | 1 | 1.984 | 0.16 |
| Habitat(P/I/rp)*Log<sub>10</sub>(PoolLength) | 2 | 15.546 | **0.0004** |
| Habitat(P/I/rp)*Log<sub>10</sub>(PoolLength)*Food | 2 | 2.618 | 0.27 |
| **Aging rate parameter** |  |  |  |
| Food | 1 | 39.967 | **<0.0001** |
| Habitat(P/I/rp) | 2 | 13.616 | **0.0011** |
| Food*Habitat(P/I/rp) | 2 | 1.943 | 0.38 |
| Log<sub>10</sub>(PoolLength) | 1 | 0.072 | 0.79 |
| Food*Log<sub>10</sub>(PoolLength) | 1 | 3.327 | 0.07 |
| Habitat(P/I/rp)*Log<sub>10</sub>(PoolLength) | 2 | 18.669 | **<0.0001** |
| Habitat(P/I/rp)*Log<sub>10</sub>(PoolLength)*Food | 2 | 0.608 | 0.74 |

Correlations between clone-specific longevity and aging parameters with the properties of habitat of origin are shown on Fig. 3 and Supplementary Fig. S3. Exclusion of 2 outliers as well as Couchy robust regression revealed a positive correlation between individual clone’s longevity parameter measured at both food levels (Fig. 3 A, B). At the same time there was no effect of habitat stability on clone’s longevity parameter (Fig. 3 C,D). There was also no effects of habitat properties on aging rate parameter, except a positive correlation between habitat size and age-specific survival measured at *ad libitum* food (Fig. S_residdeltaCSHabitats). To summarize, on the clone-level analysis clones from larger ponds tend to live longer overall and age more slowly, at least at high food level, than clones from small ponds, regardless of the stability of their habitat of origin.

**Figure 3.**
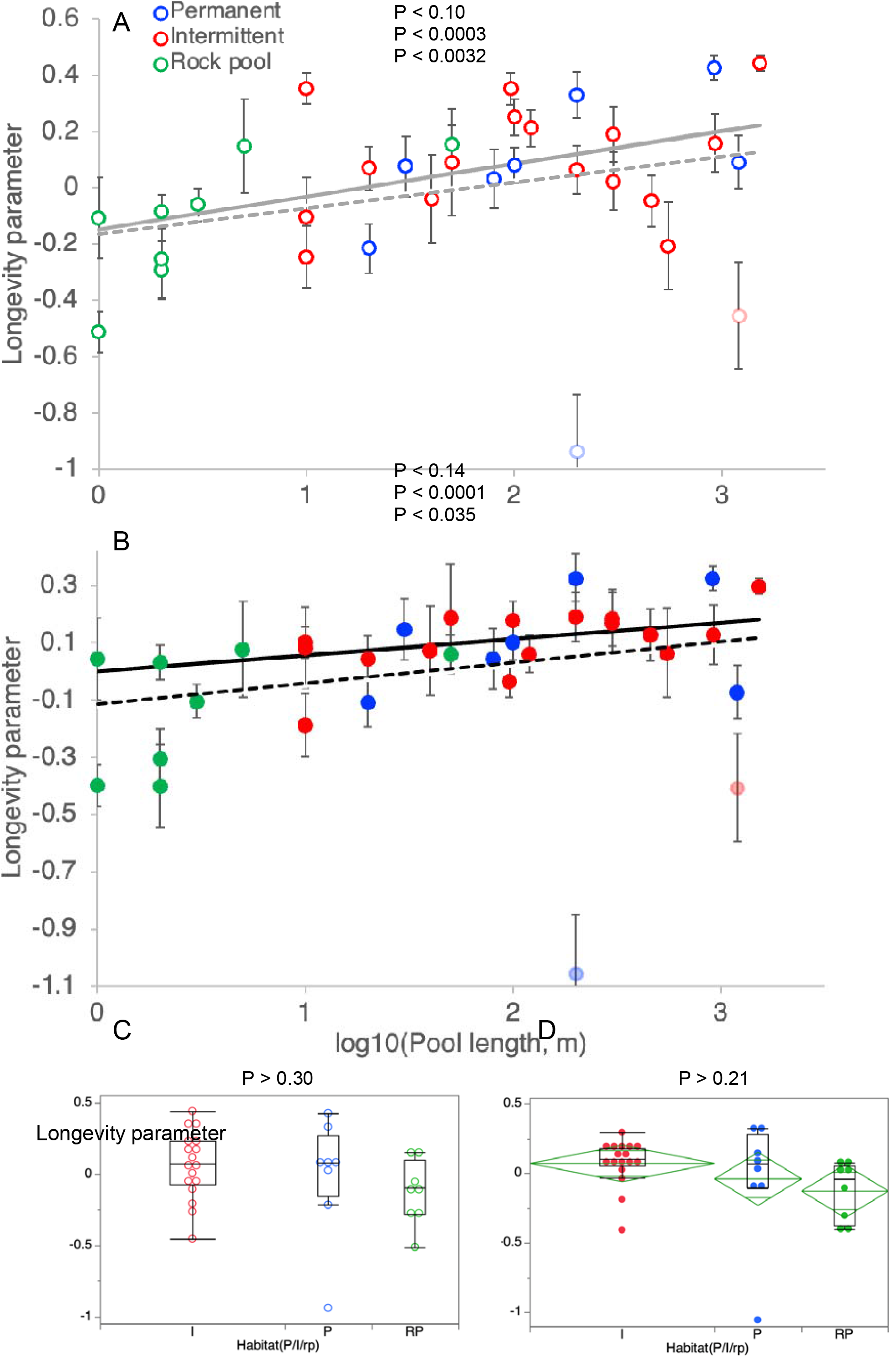
The effects of habitat of origin size (A,B) and type (Permanent, large intermittent, rock pool, C, D) on longevity parameter Weibull distribution in a Parametric survival model in 33 clones of *D.magna*. A: lifespan under restricted food. B: lifespan under *ad libitum* food. Vertical bars are SEs. Dashed lines: linear regression; solid lines: Cauchy robust regression; P-values shown are, top to bottom: MS linear regression all data; MS linear regression with two outlier clones with lifespan outside of the α = 1e-6 confidence interval (diluted colors, see also Fig. Sold3B) excluded from the analysis; and Couchy robust regression. Blue circles: clones from permanent habitats, red circles: clones from intermittent habitats, green circles: clones from rockpool intermittent habitats. Open circles: restricted food (1E5 cells/mL/day); full circles: ad libitum food (4E5 cells/mL/day).

**Fig. 4.**
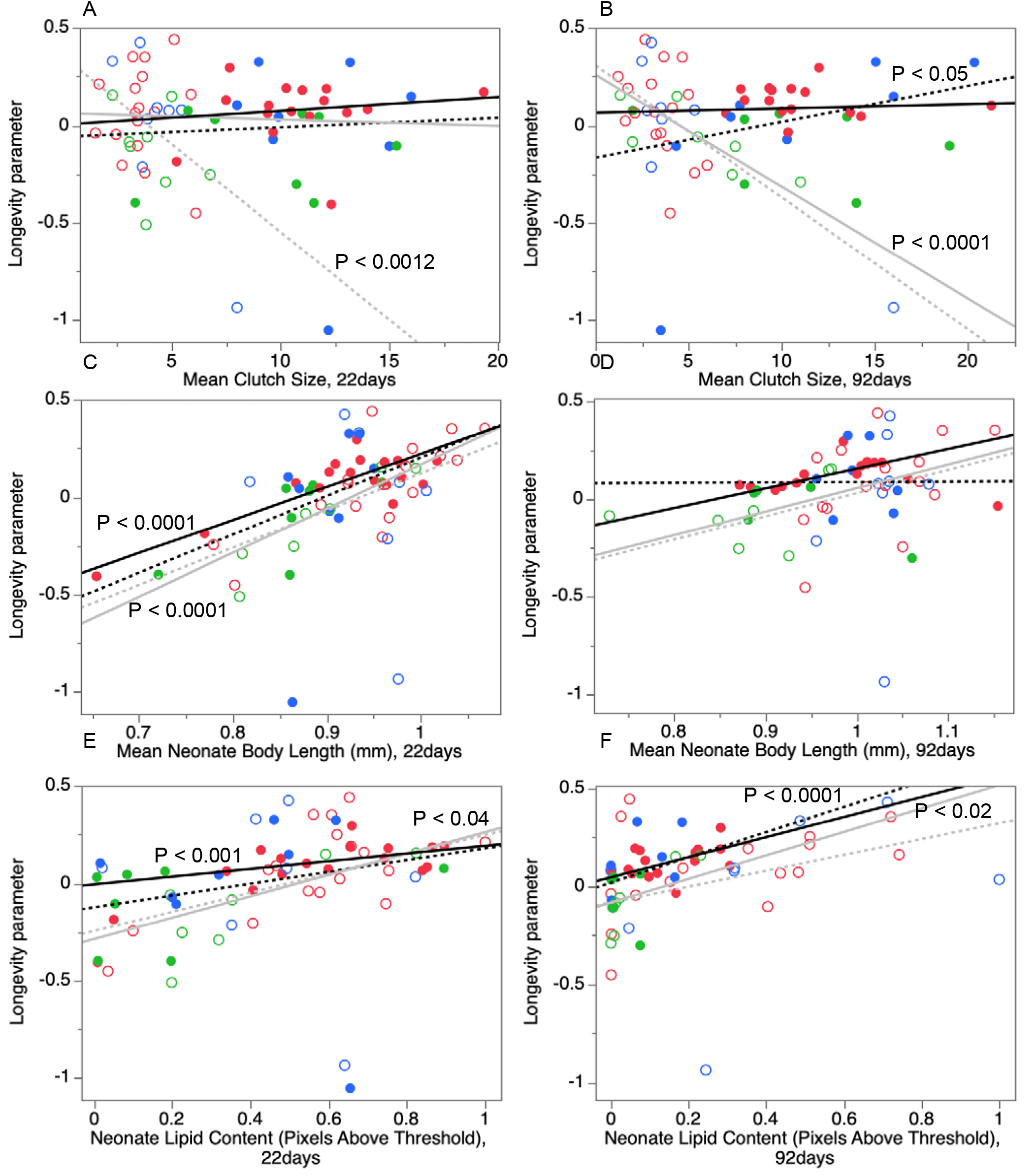
Correlations between longevity (location parameter of Weibull distribution and three component of reproductive effort, fecundity (A,B), neonate body length (C,D), and neonate lipids content (E,F) measured in young (22 days old, A, C, E) and old (92 days old, B, D, F) *Daphnia.* Symbols, lines, and colors as on Fig. 3. Only P-values <0.05 are shown.

### Longevity-fecundity trade-off

We found no evidence of trade-off between longevity and either early or late reproduction effort Fig. 3, at either food level, as most correlations between lifespan and reproductive parameters (clutch size, neonates body length, neonates lipid content) across clones were significant and positive. The only exception was the negative correlation between lifespan and fecundity (clutch size) at low food level (1e5 cells/mL/day). These correlations were significant and negative (Fig. 3A, B) both for young (22 days old) and old (92 days old) mothers. Furthermore, while in young mothers this negative correlation could be readily explained by the leverage of the two outliers with unusually low lifespan (without them regression coefficient R = -0.0103, P > 0.32), for old mothers this regression remained significant with the outliers excluded (R = -0.056, P < 0.0044). Thus, low food longevity-fecundity trade-off is not caused by the life history leverage of just a handful of clones with unusually short lifespan and high reproductive investments, but is a wide-spread phenomenon across the species, at least for the fecundity at the maternal age close to median lifespan, i.e. to the age of high mortality hazard (cf. Fig. 1). Furthermore, the negative correlations between longevity and fecundity remain significant within clones from permanent habitats considered separately (R = -0.039, P < 0.0081 and R = -0.104, P < 0.0075 for young and old mothers, respectively).

## Discussion

We successfully untangled habitat size and habitat stability as a determinant of longevity of *Daphnia* clones originating from ponds and lakes of considerable size. Longevity is higher in clones from larger habitats, regardless of habitat permanent vs. intermittent nature. It is still impossible to untangle size and stability for rock pools, which are both the smallest and the most intermittent of *Daphnia* habitats. Furthermore, we observe a positive correlation across genotypes between longevity and reproductive effort, except the one with clutch size at low food. Both observations are consistent with the genetic load explanation of intraspecific variation in lifespan (Lohr et al. 2014; Lohr & Haag 2015), but not with the adaptation to intermittent habitats that would support fecundity over longevity investment (Dudycha 2001; 2003).

Effective population size is likely to be lower in smaller ponds for other reasons than just the physical size of a water body. Smaller ponds’ *Daphnia* populations are more likely to be established from a smaller number of founders and be affected by a greater variation in egg-bank dynamics than those in larger ponds or lakes. It is likely that this is the reason that our ability to untangle habitat size and habitat stability breaks down on rock pools, as their intermittent nature makes them also most likely to contain populations of the smallest N_e_ due to repeated bottlenecks resulting from extinction/recolonization (Haag et al. 2002; Pajunen & Pajunen 2003). One may argue that adaptation to intermittent environments may occur, rather than through selection for early fecundity at the expense of longevity, through selection favoring resting eggs production – which may or may not be correlated with longevity. *Daphnia* resting (ephippial) eggs are individually more energetically expensive than parthenogenetically produced subitaneous eggs (Santos & Ebert 2022), but only two are produced each ovary cycle, resulting in potentially less severe reproduction-longevity trade-off. Yet, besides resting eggs themselves, the ephippium (resting eggs’ protective casing) implies energetic costs as well; dry mass of an ephippium with two resting eggs has been estimated as the equivalent of a clutch of 9 subitaneous eggs (Lynch 1983). Thus, whatever trade-off that might be operating between asexual reproduction and longevity would likely to be present in terms of reallocation of resources between survival and sexual reproduction.It should be noted that the neutralist explanation is the most persuasive, but not the only possible one. First, the correlation between habitat size and longevity was significant only after removal of two outliers with lifespan outside of the linear regression confidence interval corresponding to a= 1e-6, one of these outliers was a clone from a permanent and one from a large intermittent pond. This can mean that either even in relatively large zooplankton populations genotypes with heavy genetic load are possible – perhaps due to within-clone matings – or that some intermittent habitat clones do indeed persist due to investing into reproduction at the expense of longevity. The fact that the shortest-lived outlier clone also showed the highest fecundity and neonates’ lipid content (Fig. 3 A-D) is from a permanent habitat, points to the former explanation.

Second, there was clear evidence of longevity vs. offspring number trade-off ((Fig. 3A,B) at low food concentration. It means that selection does result in lower lifespan in case of high early-(Fig. 3A) or late-life (Fig. 3B) fecundity, just about equally in both permanent and intermittent habitats. However, the same effect was not observed at the high food concentration, indicating that unlimited resources permit high fecundity without a cost to longevity, to the extend allowed by each genotype’s fitness. This emphasizes that the selectionist and neutralist explanations of lifespan variation are not mutually exclusive – trade-offs may still exist and selection can still shape investments into reproduction and survival when resources are limited, on the background of different mutational loads.

While the reproduction-longevity trade-off is indeed one of the most fundamental and best documented trade-offs shaping life histories (Stearns 1992; Roff 2002; Cox et al. 2010; Lemaître et al. 2015; Ryan et al. 2024; Garratt et al. 2025), it has been long recognized that the manifestation of this trade-off is conditional on both the environment and genetic background (Hamel et al. 2009; Attisano et al. 2012; Adler et al. 2013; Cohen et al. 2020; Mc Auley 2022). First, the trade-off is ubiquitously observed across species (including *Daphnia:* Latta et al. 2011) and across diets (Attisano et al. 2012), while within a species under uniform conditions it is often overwhelmed by differences between high quality and low quality individuals (Weladji et al. 2008; Hamel et al. 2009; Coggins et al. 2021; Winder et al. 2025) leading to positive rather than negative correlations between reproduction and survival. Second, both theoretical considerations and empirical evidence point to the prediction that the reproduction-survival trade-off should only be observed when resources are limited, i.e., under caloric restriction conditions (Holliday 1989; Masoro & Austad 1996; Shanley & Kirkwood 2000; Zajitschek et al. 2019; Mc Auley 2022). This is exactly what we observed in this study: a negative correlation between longevity and reproduction (of both young and old females) is only observed at restricted food. Resource-rich conditions, thus, mask allocation trade-offs because individuals can simultaneously invest in maintenance and reproduction if their genetic background allows that; resource limitation exposes the underlying allocation conflict.

It should be noted that we did not observe a significant caloric restriction effect on longevity in majority of clones in this study, emphasizing that the effect is genotype-dependent (Beam et al 2024). We did, however, observe the extension of lifespan in each clone in which this effect was previously observed (Beam et al 2024). We explain the low frequency of clones showing caloric restriction reported here by the fact that numerous low longevity clones have been added to this study, and we previously observed that the caloric restriction lifespan extension is limited to high longevity clones (Beam et al. 2024). Yet, even in the absence of widespread shortened lifespan at *ad libitum* food level, the negative correlation between longevity and offspring number at low food persists. This indicates that high reproduction investments result in shorter lifespan across genotypes, but not necessarily across environments.

Furthermore, the reproduction-longevity correlation is negative only for offspring number, but not offspring quality (body size and lipid content). This indicates that the observed effect is caused by genotype-specific adaptive re-allocation rather than overall shortage of resources (cf. Piper et al. 2023) – which would have affected both offspring number and quality. It has been repeatedly demonstrated that *Daphnia* produce fewer but larger and better provisioned offspring when resources are limited (Boersma 1995; McKee & Ebert, 1996).

## Conclusions

Intraspecific variation in longevity in *Daphnia* is better explained by the size of the habitat of origin, as a proxy of population size, than by habitat stability, consistent with genetic load as the reason behind shorter lifespans. Reproduction-longevity tradeoff is observed only at low food and only between longevity and offspring number, but not between longevity and offspring quality. This supports neutralist drift- and deleterious mutations-based view of intraspecific variation in lifespan and early reproduction.

## Acknowledgements

We are grateful to Jeff Dudycha and Christoph Haag for useful discussions, to Dieter Ebert for providing and Piet Schpaak for transporting *Daphnia* clones, and to Meridith Smith and Jacob Halfacre for laboratory assistance.

## Supplementary materials

### Supplementary Tables

**Table S1.**
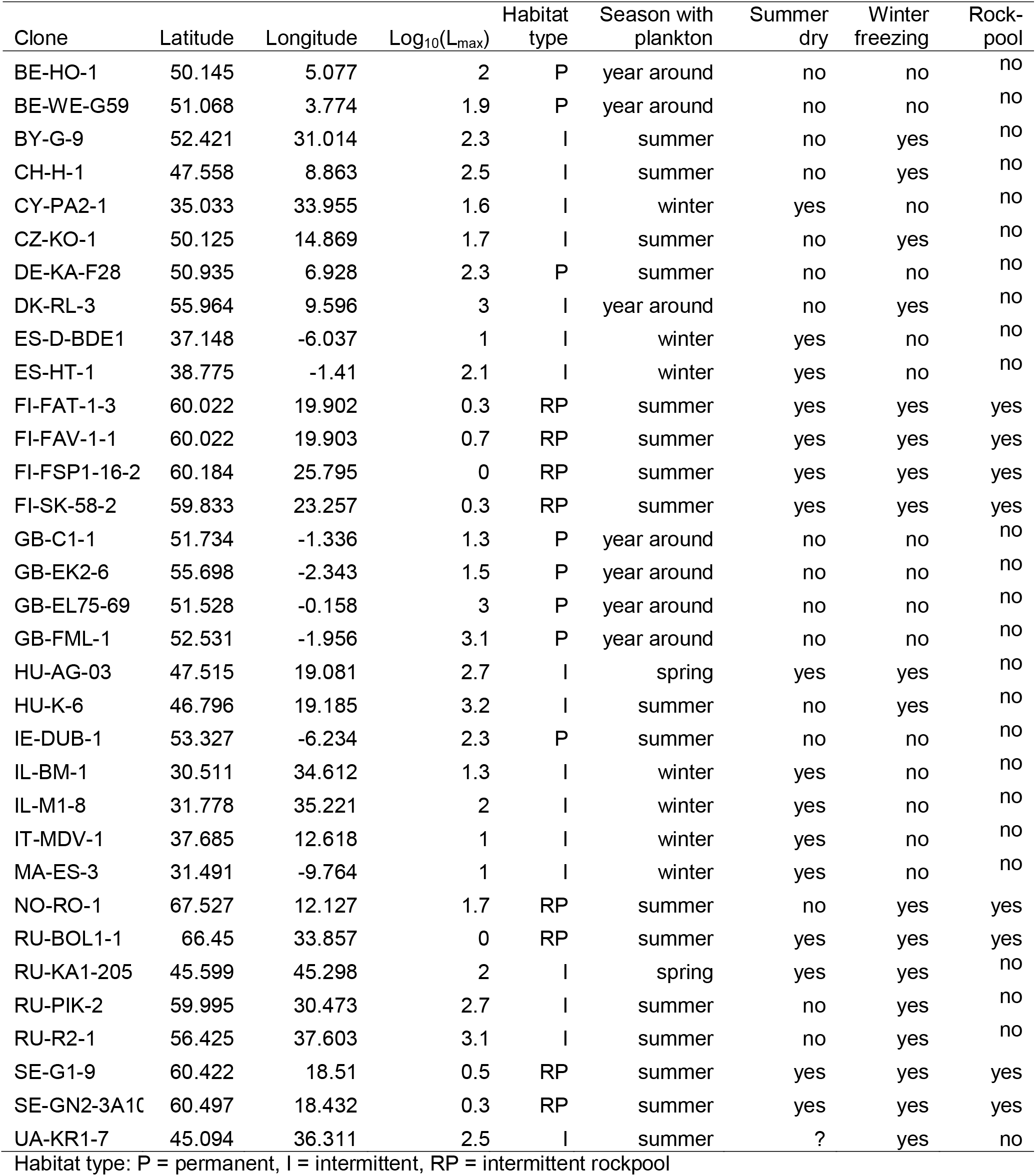
Location, water body size, and habitat stability of clones’ origin (D.Ebert, personal communication).

| Clone | Latitude | Longitude | Log<sub>10</sub>(L<sub>max</sub>) | Habitat type | Season with plankton | Summer dry | Winter freezing | Rock-pool |
|---|---|---|---|---|---|---|---|---|
| BE-HO-1 | 50.145 | 5.077 | 2 | P | year around | no | no | no |
| BE-WE-G59 | 51.068 | 3.774 | 1.9 | P | year around | no | no | no |
| BY-G-9 | 52.421 | 31.014 | 2.3 | I | summer | no | yes | no |
| CH-H-1 | 47.558 | 8.863 | 2.5 | I | summer | no | yes | no |
| CY-PA2-1 | 35.033 | 33.955 | 1.6 | I | winter | yes | no | no |
| CZ-KO-1 | 50.125 | 14.869 | 1.7 | I | summer | no | yes | no |
| DE-KA-F28 | 50.935 | 6.928 | 2.3 | P | summer | no | no | no |
| DK-RL-3 | 55.964 | 9.596 | 3 | I | year around | no | yes | no |
| ES-D-BDE1 | 37.148 | -6.037 | 1 | I | winter | yes | no | no |
| ES-HT-1 | 38.775 | -1.41 | 2.1 | I | winter | yes | no | no |
| FI-FAT-1-3 | 60.022 | 19.902 | 0.3 | RP | summer | yes | yes | yes |
| FI-FAV-1-1 | 60.022 | 19.903 | 0.7 | RP | summer | yes | yes | yes |
| FI-FSP1-16-2 | 60.184 | 25.795 | 0 | RP | summer | yes | yes | yes |
| FI-SK-58-2 | 59.833 | 23.257 | 0.3 | RP | summer | yes | yes | yes |
| GB-C1-1 | 51.734 | -1.336 | 1.3 | P | year around | no | no | no |
| GB-EK2-6 | 55.698 | -2.343 | 1.5 | P | year around | no | no | no |
| GB-EL75-69 | 51.528 | -0.158 | 3 | P | year around | no | no | no |
| GB-FML-1 | 52.531 | -1.956 | 3.1 | P | year around | no | no | no |
| HU-AG-03 | 47.515 | 19.081 | 2.7 | I | spring | yes | yes | no |
| HU-K-6 | 46.796 | 19.185 | 3.2 | I | summer | no | yes | no |
| IE-DUB-1 | 53.327 | -6.234 | 2.3 | P | summer | no | no | no |
| IL-BM-1 | 30.511 | 34.612 | 1.3 | I | winter | yes | no | no |
| IL-M1-8 | 31.778 | 35.221 | 2 | I | winter | yes | no | no |
| IT-MDV-1 | 37.685 | 12.618 | 1 | I | winter | yes | no | no |
| MA-ES-3 | 31.491 | -9.764 | 1 | I | winter | yes | no | no |
| NO-RO-1 | 67.527 | 12.127 | 1.7 | RP | summer | no | yes | yes |
| RU-BOL1-1 | 66.45 | 33.857 | 0 | RP | summer | yes | yes | yes |
| RU-KA1-205 | 45.599 | 45.298 | 2 | I | spring | yes | yes | no |
| RU-PIK-2 | 59.995 | 30.473 | 2.7 | I | summer | no | yes | no |
| RU-R2-1 | 56.425 | 37.603 | 3.1 | I | summer | no | yes | no |
| SE-G1-9 | 60.422 | 18.51 | 0.5 | RP | summer | yes | yes | yes |
| SE-GN2-3A1[illegible] | 60.497 | 18.432 | 0.3 | RP | summer | yes | yes | yes |
| UA-KR1-7 | 45.094 | 36.311 | 2.5 | I | summer | ? | yes | no |

Habitat type: P = permanent, I = intermittent, RP = intermittent rockpool

**Supplementary Table S2.**
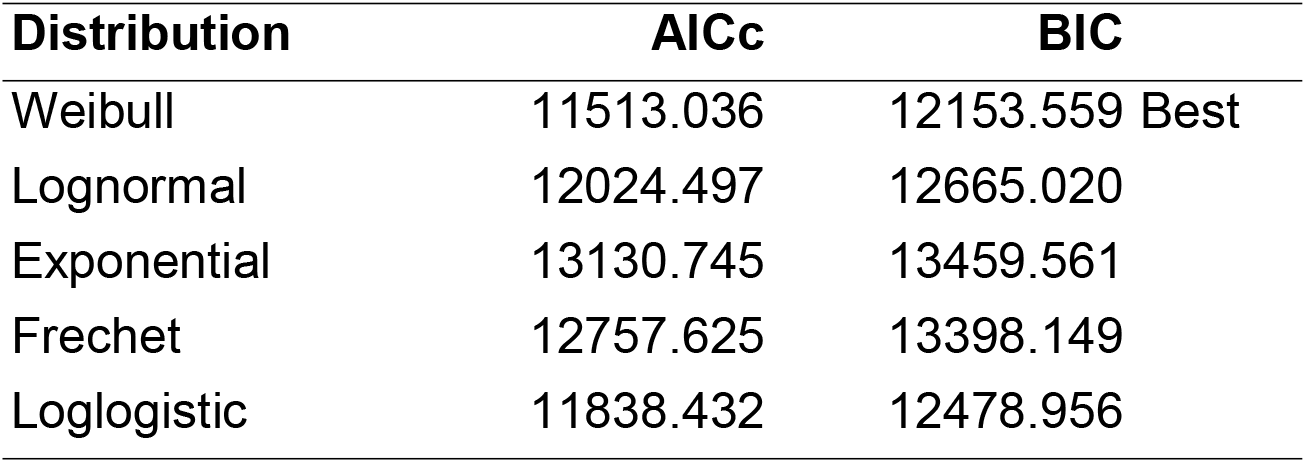
Distributions comparison for the Parametric Survival Model.

**Supplementary Table S3.** Results of Royston–Parmar flexible parametric survival model, implemented in R with rstpm2::stpm2().

| Factor | Estimate | Std. Error | z value | Pr(z) |
|---|---|---|---|---|
| Intercept | -3.240 | 0.404 | -8.025 | 1.01e-15 |
| Natural spline of log(age), df=2, 1 | 1.685 | 0.831 | 2.027 | 0.043 |
| Natural spline of log(age), df=2, 2 | 2.889 | 0.416 | 6.939 | 3.9e-12 |
| Habitat_I | -0.635 | 0.484 | -1.310 | 0.19 |
| Habitat_RP | -2.155 | 0.930 | -2.317 | 0.020 |
| Food_400 | -2.633 | 0.397 | -6.633 | 3.3e-11 |
| Log<sub>10</sub>(PoolLength) | -0.463 | 0.460 | -1.006 | 0.31 |
| Habitat_I : Food_400 | 0.115 | 0.170 | 0.677 | 0.50 |
| Habitat_RP : Food_400 | 0.461 | 0.341 | 1.353 | 0.17 |
| Habitat_I : Log<sub>10</sub>(PoolLength) | 0.585 | 0.443 | 1.319 | 0.19 |
| Habitat_RP : Log<sub>10</sub>(PoolLength) | -0.461 | 0.568 | -0.812 | 0.42 |
| Food_400 : Log<sub>10</sub>(PoolLength) | 0.222 | 0.196 | 1.131 | 0.26 |
| Habitat_I : log(age) | 0.216 | 0.526 | 0.411 | 0.68 |
| Habitat_RP : log(age) | 1.770 | 0.952 | 1.859 | 0.063 |
| Log<sub>10</sub>(PoolLength : log(age) | 0.007 | 0.344 | 0.021 | 0.98 |
| Food_400 : log(age) | 3.654 | 0.547 | 6.679 | 2.4e-11 |
| Habitat_I : Food_400 : Log<sub>10</sub>(PoolLength) | -0.342 | 0.223 | -1.536 | 0.12 |
| Habitat_RP : Food_400 : Log<sub>10</sub>(PoolLength) | -0.104 | 0.287 | -0.363 | 0.72 |
| Logtheta (Clone-level frailty variance) | -0.999 | 0.262 | -3.810 | 0.00014 |

**Table S4.** Proportional Hazards (Wald test) for the effects of clones, food level and their interaction on longevity.

| Source | DF | Wald ChiSquare | Prob>ChiSq |
|---|---|---|---|
| Food | 1 | 5.045 | 0.025 |
| Clone | 32 | 513.133 | <0.0001 |
| Food*Clone | 32 | 50.969 | 0.018 |

### Supplementary figures

**Figure S1.**
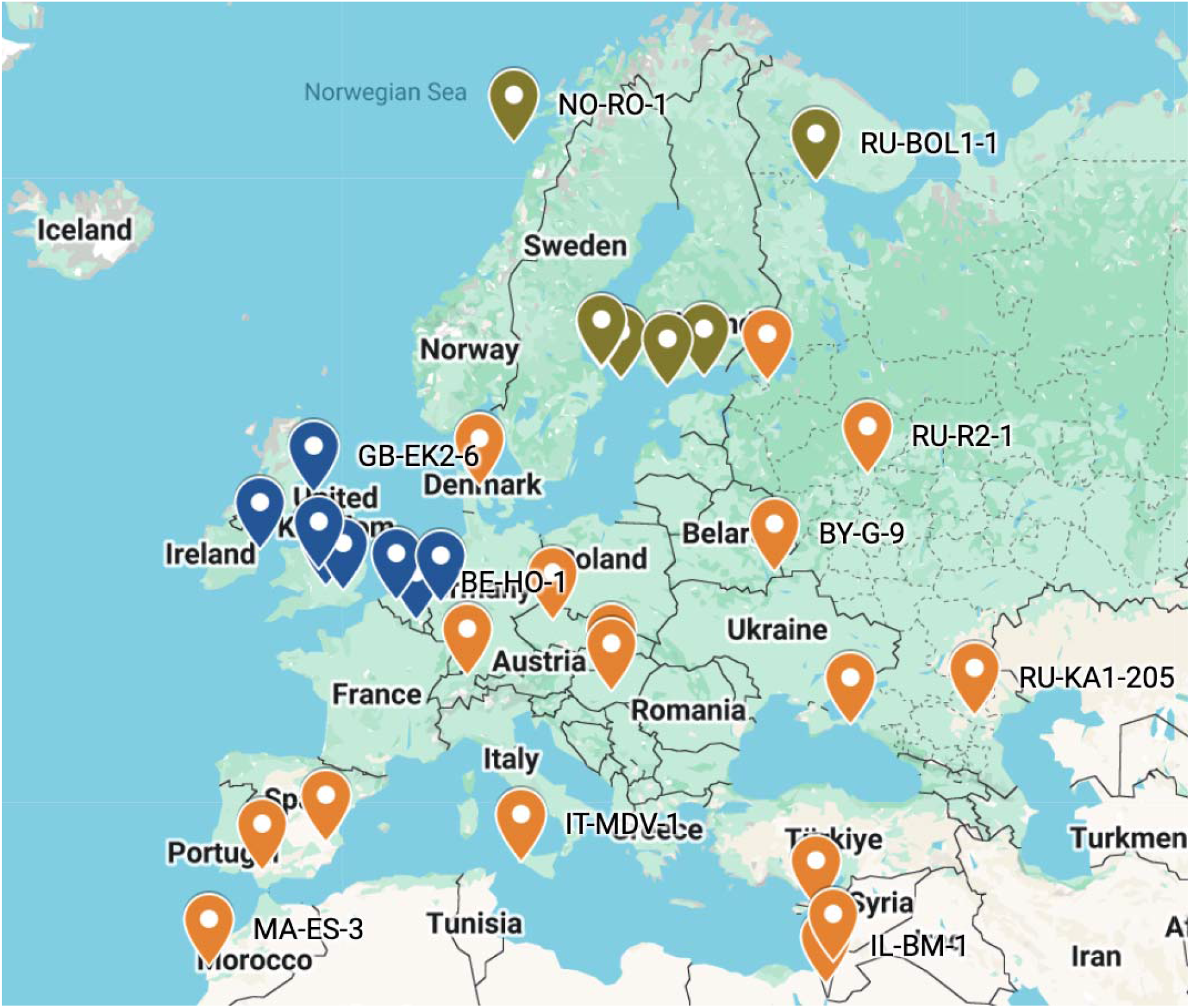
Locations of *D. magna* clones’ origins. Orange – intermittent habitats, blue – permanent habitats, dark green – rockpools. Select clone’s locations labeled. See Table S1 for detailed location information.

**Figure S2.**
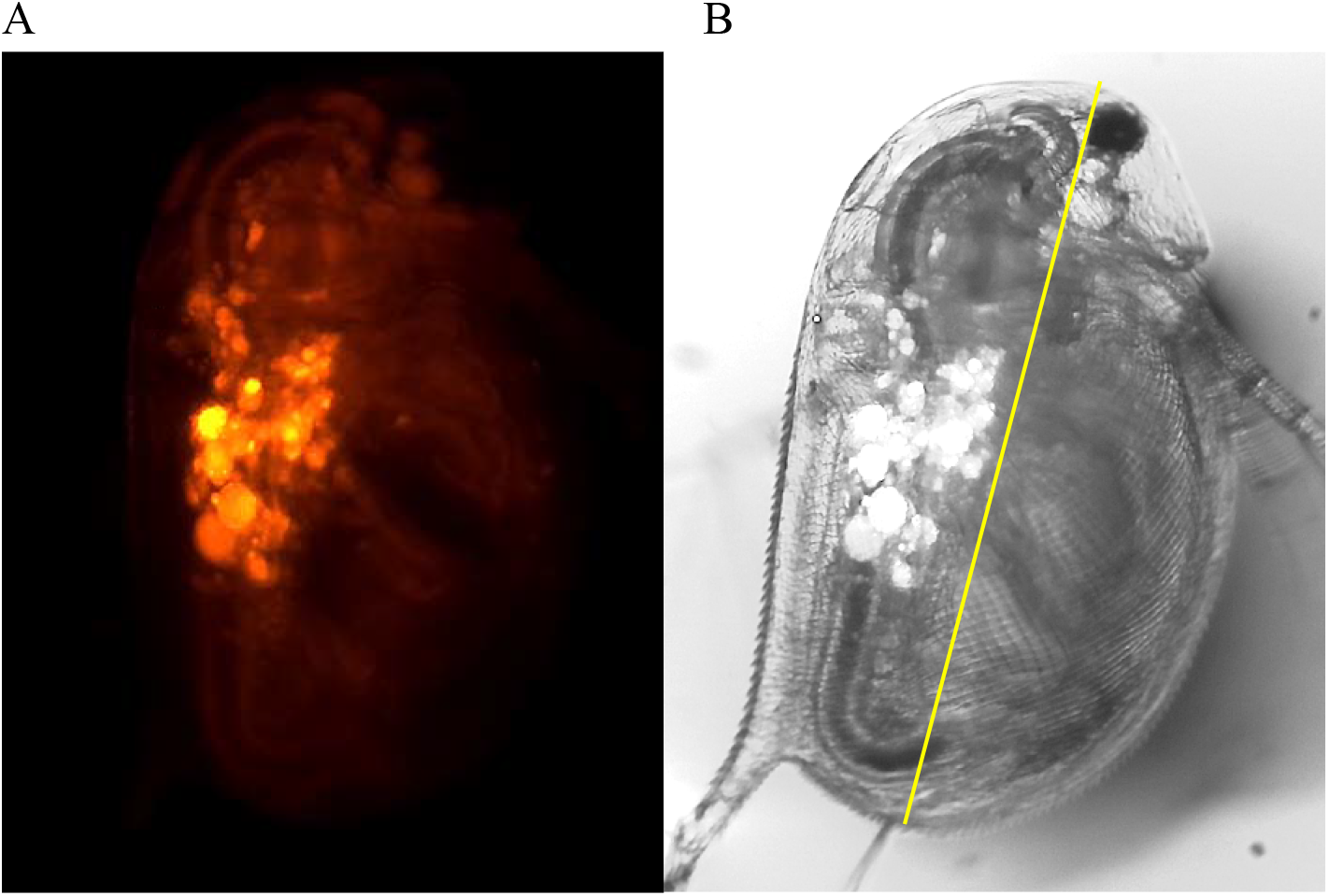
Examples of neonate images (same individual) for lipids quantification (Nile Red stain, green excitation, red emission, A) and body length measurement (brightfield, B).

**Supplementary Fig. S3.**
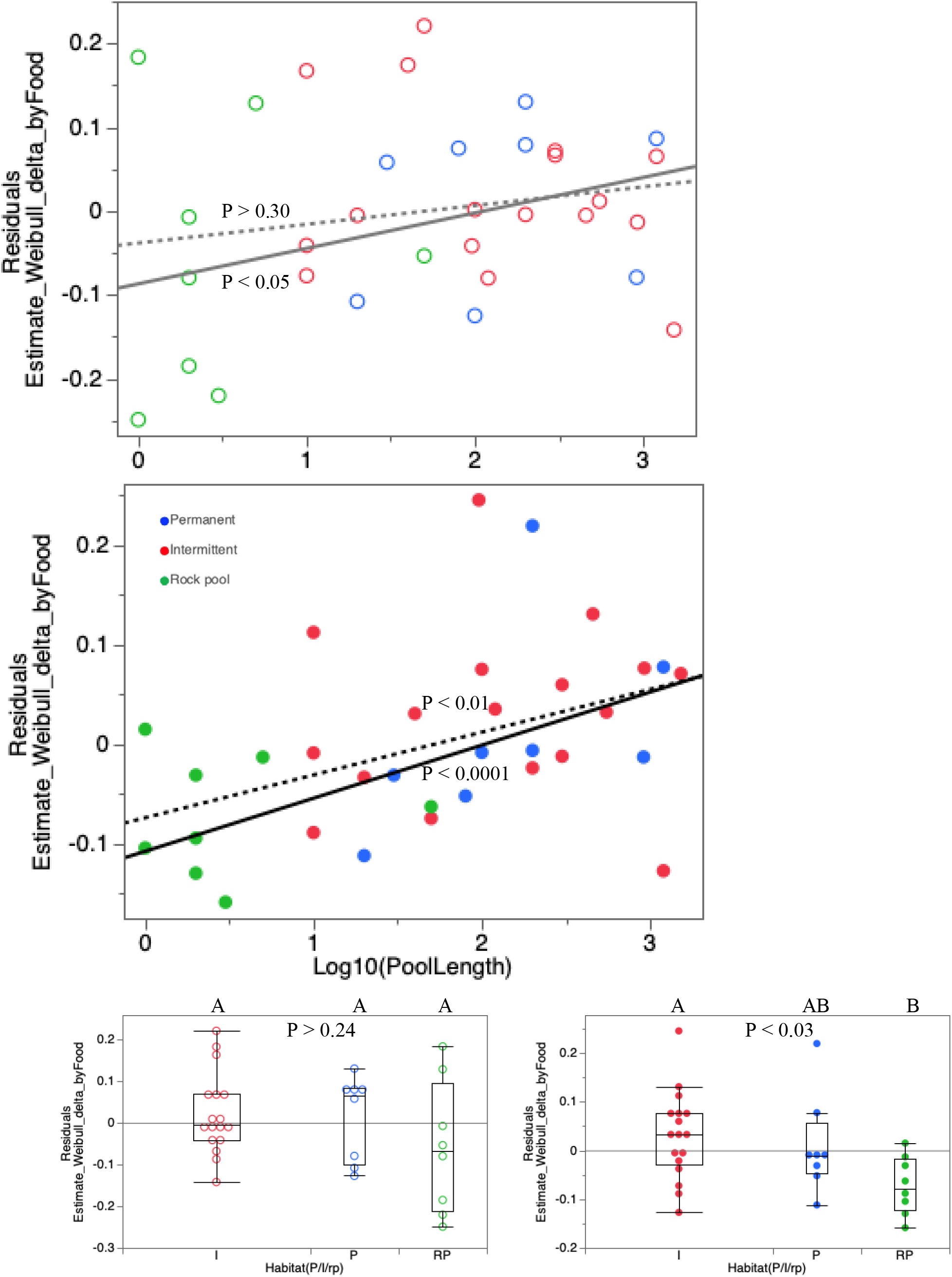
Correlation between clone- and food level-specific aging parameter delta (residuals from linear regression over longevity parameter) with habitat size (A, B) and stability (C,D). Symbols as on Main text Fig. 3.

